# Novel mosaic mice with diverse applications

**DOI:** 10.1101/2020.03.21.001388

**Authors:** Yuxin Chen, Shaoshuai Mao, Bo Liu, Zhengyu Jing, Ying Zang, Jing Xia, Jianlong Sun, Tian Chi

## Abstract

Gene-deficient mouse models are indispensable for interrogating mammalian gene functions, but the conventional models allow the study of only one or few genes per mouse line, which has been a bottleneck in functional genomics. To confront the challenge, we have combined the CRISPR-Cas and Cre-Lox systems to develop a novel type of mosaic mice termed **MARC** (**M**osaic **A**nimal based on g**R**NA and **C**re) for targeting many genes per mouse but only one gene per cell. This technology employs a transgene comprising a modified U6 promoter upstream of a series of floxed gRNA genes linked together in tandem, with one gRNA expressed per cell following Cre-mediated recombination. At least 61 gRNA genes can be stably maintained in the transgene, and importantly, enables robust proof-of-principle *in vivo* screens, demonstrating the potential for quickly evaluating the functions of many genes in diverse tissues in a single MARC line. In theory, MARC can also be analyzed by single-cell sequencing, and should enable cost-effective derivation of conventional single-gene-KO lines via simple breeding. Our study establishes MARC as an important addition to the mouse genetics toolbox.

## Introduction

In the post-genomic era, the genomes of diverse organisms have been sequenced, but their functions remain largely a mystery. Gene targeting in mice is the gold standard for uncovering gene functions in mammals and for modeling human hereditary diseases (Bouabe and Okkenhaug, 2013; Capecchi, 2005; Nguyen and Xu, 2008). It involves, in the simplest form, knockout of a single gene either globally or in cell-type specific manner. As only one target gene is studied per mouse line, the throughput is very low, which hinders functional genomics research.

A potential strategy to overcome the limitation is to use a genetic mosaic model where many different genes are collectively knocked out in a single mouse, but only one gene per cell. Such mice may lack obvious organismal phenotypes even when every single cell in the body is targeted, as the KO may be inconsequential in a large fraction of the cells in the body. This is a limitation of the mosaic mice, but also an advantage, in that it enables the study of cell-intrinsic defects reflecting the direct effect of the KO within the cells, without the confounding influences secondary to the KOs in other cells (Nguyen and Xu, 2008; Rossant and Spence, 1998). The mosaic mice may be used in several ways. If the germ cells are targeted, the mosaic founders could obviously be used to produce many different lines of single-gene KO offspring via simple breeding, thus reducing the cost of making KO lines. Most excitingly, many genes could be directly studied in a single mosaic mouse using forward genetic screens.

Forward genetic screens entail the introduction of mutations at many genes, followed by selection of cells with desired phenotypes and characterization of their associated mutations, thus linking phenotype to genotype (Shalem et al., 2015). Galvanized in recent years with the introduction of the revolutionary CRISPR-Cas technology, forward genetic screens have proven a powerful strategy for discovering mammalian gene functions (Doench, 2018; Joung et al., 2017; Shalem et al., 2015). However, the screens have been done mainly in cultured cells, which severely limits its impact, given that many important physiological and pathological processes cannot be (faithfully) recapitulated *in vitro*.

Multiple strategies at genetic screens in live mice have been reported. First, genetic screens have been achieved in mosaic mice generated via transposon-mediated insertional mutagenesis (Copeland and Jenkins, 2010; Friedrich et al., 2019; Rad et al., 2010). As the insertions are monoallelic and occur mostly at the vast intronic regions in the genome, the overwhelming majority of the insertional events are inconsequential, and so the mosaic mice have been successfully used only in the screens for cancer genes, where even a very rare phenotype can become detectable thanks to the signal amplification resulting from cancer growth.

A broader range of phenotypes in mice can be screened using pooled viral gRNA or shRNA libraries focused on the protein-coding genes, but the screens can be performed only in special cells. Specifically, two forms of screens using pooled libraries have been documented. In the first, mouse cells are isolated, transduced with the gRNA libraries and adoptively transferred into recipients where they are screened (Dong et al., 2019; LaFleur et al., 2019). This method is applicable only to transplantable cells such as lymphocytes and bone marrow progenitor cells. Furthermore, isolation and *ex vivo* manipulations of the isolated cells are technically demanding, and the procedures may also change the cellular properties. The second form of *in vivo* screens is done *in situ*, with the cells residing in their native environment and the viral libraries delivered via injection. *In situ* screens bypass *ex vivo* manipulations, but is doable only in a few accessible cell types, primarily the epithelial cells (the lung epithelium and embryonic epidermis) and hepatocytes(Beronja et al., 2013; Laurin et al., 2019; Rogers et al., 2018; Schramek et al., 2014; Wang et al., 2018; Winters et al., 2018). Furthermore, the library deliveries require skills and sometimes special instruments. For example, the genetic screen in the embryonic epidermis requires ultrasound-guided injection of the viral library into the amniotic fluid *in utero* (Beronja et al., 2013; Laurin et al., 2019; Schramek et al., 2014).

We now describe a novel type of mosaic mice with multiple potential applications, including dramatically simplified *in vivo* screening that is free from the limitations of the existing screening methods mentioned above.

## Results

### Experimental Design

At the heart of MARC is a transgene comprising three parts (Fig. 1A). The first is a modified U6 promoter carrying a bifunctional LoxP site (TATA-lox71), which is not only able to undergo Cre-mediated recombination, but also contains a functional TATA box in its spacer region to direct gRNA expression(Ventura et al., 2004). The second part of the transgene is a gRNA gene immediately downstream of the U6 promoter; this position is termed Position 0 (P0). The gRNA gene ends at a transcription terminator (a string of Ts). Placed downstream of the P0 gRNA gene is the final component of the transgene, which is a series of tandemly linked gRNA genes (#1, 2, 3…), each ending at a transcription terminator and floxed with another bifunctional LoxP site (TATA-loxKR3). Lox71 and LoxKR3 are LoxP variants bearing mutations at the 5’ and 3’ inverted repeats, respectively; following recombination between the two variants, the resulting LoxP site would carry mutations at both ends, preventing its further recombination(Araki et al., 2010). In the transgenic mice, the P0 gRNA gene is constitutively expressed, while a gRNA downstream can be induced whenever Cre-mediated recombination takes place between Lox71 at the U6 promoter and LoxKR3 at the 5’ end of the particular gRNA gene; such recombination would remove the intervening sequence between the U6 promoter and the gRNA gene, thus relocating the gRNA gene to P0 for expression (Fig. 1B). As discussed before, the U6 promoter is supposed to undergo recombination only once, thus avoiding sequential expression of different gRNAs and hence the potential targeting of multiple genes within the same cells. Of note, the physical location of the gRNA genes along the transgene can affect the probability of recombination and hence of gRNA expression, but in general, all gRNAs in the transgene can be expressed and the corresponding target genes knocked out in the presence of Cas9, creating mosaic mice with multiple potential applications including genetic screening, single-cell RNA-Seq and derivation of single-gene KO lines (Fig. 1C).

**Fig. 1.**
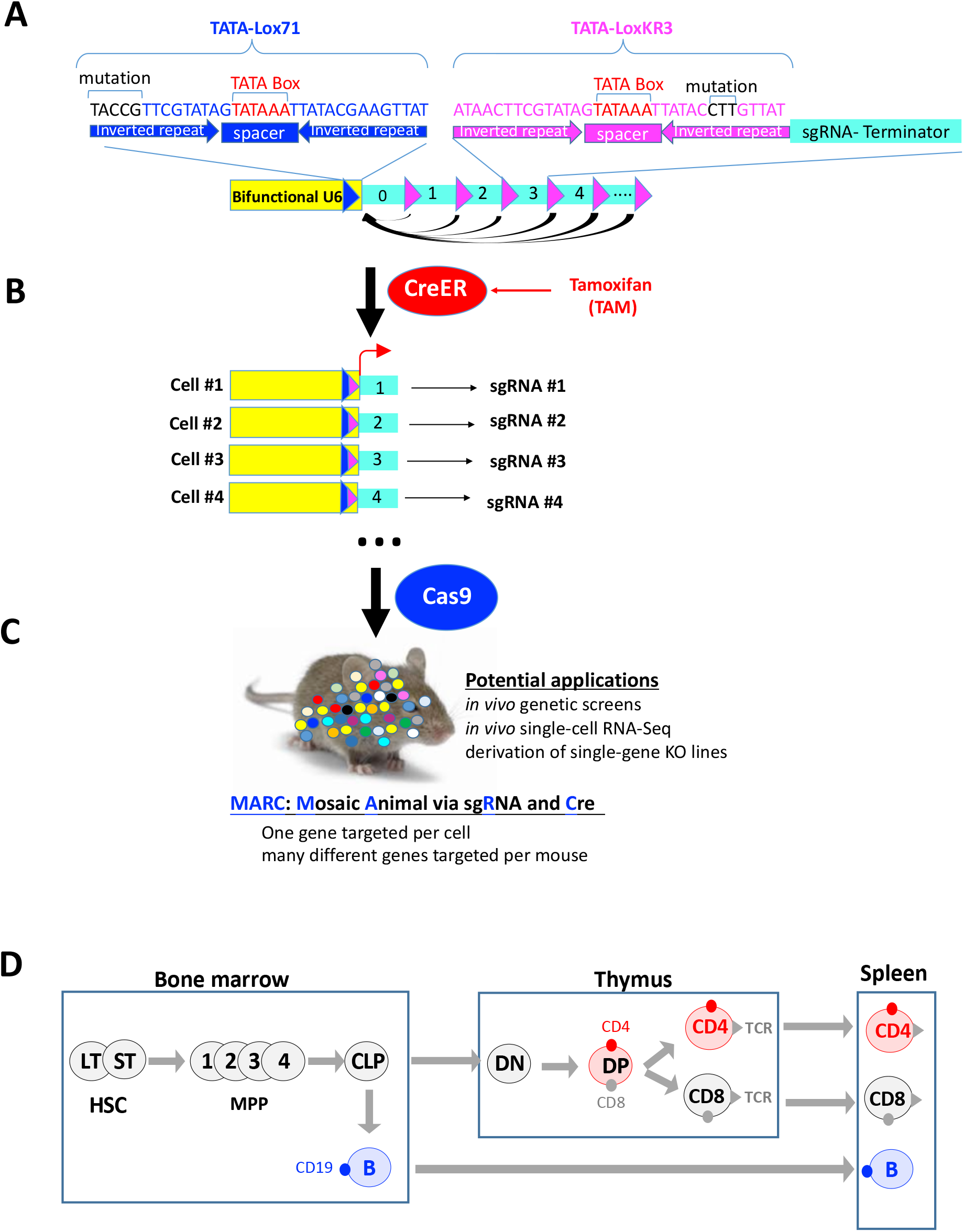
Principle of MARC (A-C) and the major experimental system for its evaluation (D) A) The transgene before recombination. The cluster of arrows below the transgene denote the “productive” recombination events that lead to the expression of various gRNAs. Recombination can also occur among the floxed gRNA genes, which shortens the transgene and may indirectly affect the productive recombination (not shown). The mutations in the two LoxP variants (Lox71 and LoxKR3) are indicated with black letters. B) The recombined transgene. In cells expressing CreER, tamoxifen (TAM) treatment activates Cre, which then catalyzes the recombination to induce transgene expression in a cell-specific manner. The recombined transgenes may retain a single gRNA gene (depicted), multiple gRNA genes (with the first gene immediately downstream of the U6 promoter expressed; not shown) or none gRNA gene (not shown), the latter produced when Lox71 is recombined with the last LoxKR3 (which is engineered in an attempt to reduce the bias in gRNA representation after recombination). In this study, ubiquitously expressed CreER (from the Ubc promoter) is used to catalyze the recombination in diverse tissues (Ruzankina et al., 2007). C) Mosaic mice are generated using the expressed gRNAs and Cas9. In this study, Cas9 is ubiquitously expressed from the CAG promoter for gene KO in diverse tissues(Chu et al., 2016a, 2016b). D) Simplistic view of hematopoiesis. HSC, hematopoietic stem cell, which has two subsets: long-term (LT) and short-term (ST); MPP, multipotent progenitor, which has 4 subsets: MPP1-4; CLP, common lymphoid progenitor; MyP, myeloid progenitor; DN, Double negative (CD4^-^D8^-^); DP, Double Positive (CD4^+^CD8^+^). CD4 (red dot) is expressed in DP and CD4 T cells as depicted, whereas CD47 is constitutively expressed throughout T cell development (not shown)

The success of the MARC technology is contingent on multiple conditions: the repetitive transgene must be stably maintained *in vivo*, Cre-mediated recombination must be efficient enough to relocate different gRNA genes on the array to P0, and finally, once relocated, the P0 gRNA must be efficiently expressed. The MARC concept is obviously risky, particularly because repetitive sequences are prone to deletion and epigenetic silencing.

To test the MARC concept, we expressed CreER and Cas9 ubiquitously (from the Ubc and CAG promoters, respectively) (Chu et al., 2016a, 2016b; Ruzankina et al., 2007), so as to examine MARC performance in diverse organs. However, our focus is on the hematopoietic cells, as they are readily quantifiable using FACS (Fig. 1D). Specifically, we focused on three well-defined surface markers: CD19, CD4 and CD47, which are expressed on B cells, some T cells and diverse cell types (including T cells), respectively. Importantly, the elimination of these surface markers does not affect cell proliferation or survival under our assay conditions, thus facilitating the analysis (Guimont-Desrochers et al., 2009; Rahemtulla et al., 1991; Rickert et al., 1995).

### An 11-gRNA transgene inserted into the *H11* locus: the first proof of the MARC concept

We started the project by generating a mouse line carrying a relatively short transgene comprising 11 gRNA genes targeting the three aforementioned markers: CD19, CD4 and CD47 (Fig. 2A, top). CD19 is targeted by a single gRNA, whereas CD4 and CD47 each by 5 distinct gRNAs. The CD19 gRNA gene is placed at P0, thus blocking the expression of other gRNAs until Cre-mediated recombination. In addition, the CD19 gRNA is predicted to be constitutively expressed starting in embryos, which should lead to maximal levels of knock-out, thus serving as a positive control for CD4 and CD47 targeting initiated in adults. The 11-gRNA transgene was inserted into the safe harbor *H11* locus (Hippenmeyer et al., 2010)(Fig. 2A, bottom; Fig. 2B), which was selected instead of the popular *Rosa26* locus because the latter has been occupied by the CAG-Cas9 transgene. We bred and analyzed the *11-gRNA array; Ubc-CreER; CAG-Cas9* mice (termed MARC-11), where the gRNA, Cas9 and CreER were all widely expressed. We found that in adult mice, CD19 expression was abolished in ~70% of splenic B cells, with 30% of B cells retaining CD19 expression, indicating the KO was incomplete (Fig. 2C, left). To confirm this, we isolated the genomic DNA from the splenocytes, PCR-amplified the gRNA target site at the *CD19* locus and quantified the indels using Sanger sequencing. In general agreement with the FACS analysis, a sizable fraction (31%) of the genomic DNA was unedited, with the remaining edited alleles displaying highly heterogeneous indels, indicating that the KO was rather inefficient and occurred late during development (Fig. 2C, heatmap, middle). Editing was more efficient in the thymus (90%) but not the kidney, liver and brain (30%- 50%; Fig. 2C, heatmap, middle). The KO efficiencies in different organs tended to vary with the CD19 gRNA expression levels, which peaked in the thymus (Fig. 2C, right). However, even in the thymus, the gRNA expression reached only ~6% of the *Actinb* level, suggesting there was much room for improvement of gRNA expression.

**Fig. 2.**
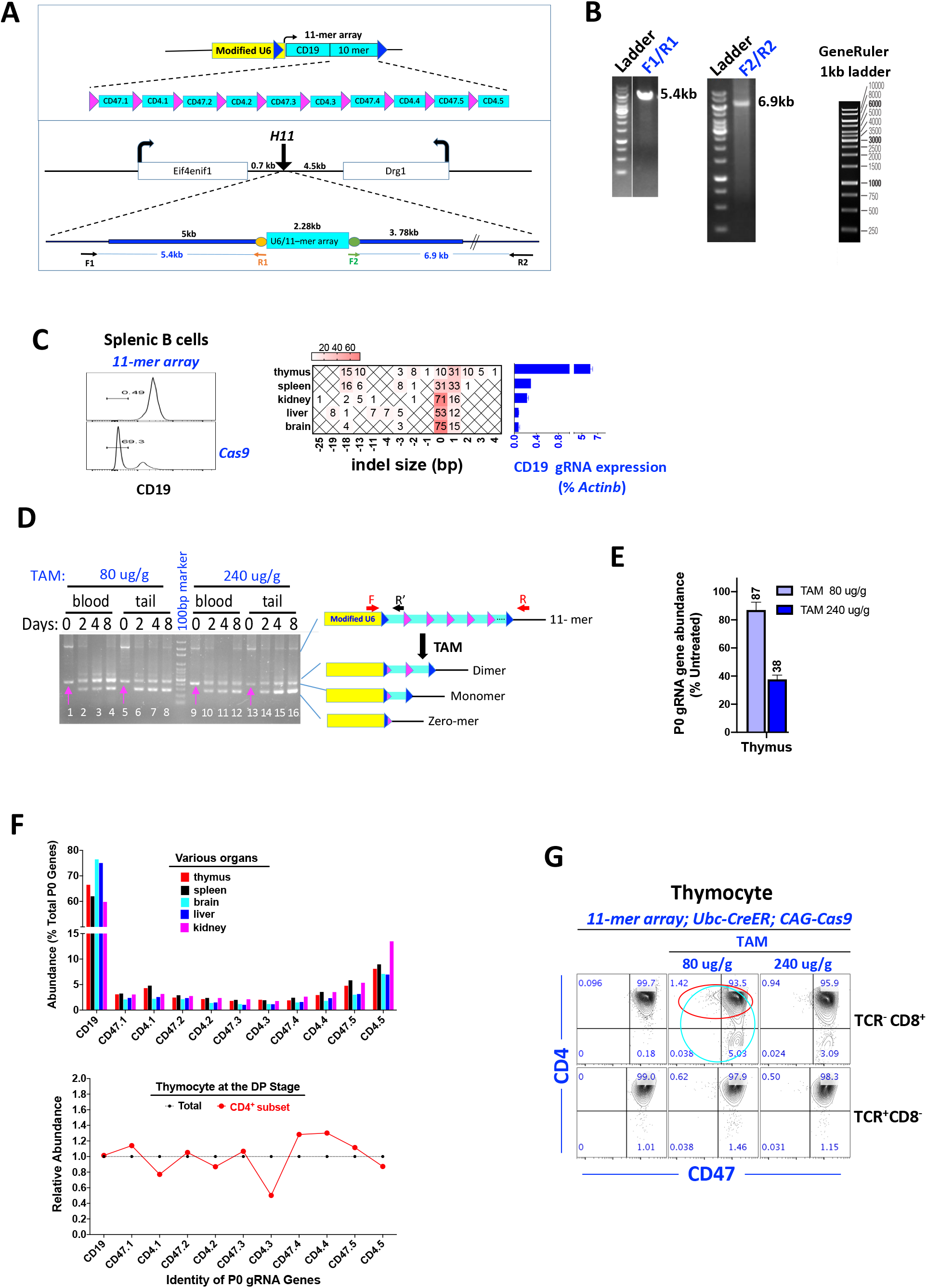
Creation and characterization of MARC-11. A-B) Generation of MARC-11. The 11-gRNA array contains a single CD19 gRNA gene, 5 CD47 gRNA genes (CD47.1 to CD47.5) and 5 CD4 gRNA genes (CD4.1 to CD4.5; Fig. 2A, top). The transgene was inserted into the *H11* locus (Fig. 2A, bottom). The thick blue lines flanking the transgene are the homology arms in the targeting vector. F1/R1 and F2/R2 are PCR primers for screening the founders (Fig. 2B). C) Quantification of CD19 expression by FACS (left), of indels at the CD19 locus by PCR-Sanger sequencing (middle) and of the CD19 gRNA expression by qRT-PCR (right). Values from the qRT-PCR are mean+/-SD. D) Transgene recombination as revealed by PCR detecting all forms of transgenes including intact transgene and “zero-mer”. PCR primer pair is F/R (F/R’ was used instead for experiments in Fig. 2E). The pink arrows indicate a nonspecific amplicon that only appeared in the absence of recombination and migrated slightly faster than the monomer. For unknown reasons, residual amounts of intact array tended to persist even when dimers were fully undetectable. Samples were from F6 mice (counting the founders as F1). E) Total abundance of various P0 gRNA genes as a whole in TAM treated mice relative to that in untreated mice. Thymocytes were analyzed on Day 8 following TAM. The primer pair used (F/R’) are depicted in Fig. 2D, with the R’ binding the scaffold common to all gRNA genes. The qPCR signals were normalized to an internal control (*Brg1*) and plotted relative to that of the samples lacking TAM treatment. F) Representation of individual gRNA genes at P0 following recombination. Samples were harvested on Day 8 following TAM (80 ug/g). PCR was performed using a primer pair located at similar positions to F/R’ depicted in Fig. 2D (F3/R3, not shown), and the P0 gRNA gene identified by NGS. The representation of gRNA genes at P0 in various organs was shown at the top, whereas the bottom plot shows the P0 gRNA gene representation in the CD4^+^ subset of the TCR^-^CD8^+^thymocytes (TCR^-^D4^+^CD8^+^) relative to the unfractionated counterpart (TCR-CD8^+^; see Fig. 2G for the two populations). G) FACS analysis of CD4 and CD47 expression in thymocytes. Cells were stained with CD4, CD8, TCR and CD47 antibodies before the subsets that should normally express CD4 (namely, the cells at the DP and CD4 cell stages, marked by TCR^-^D8^+^ and TCR+CD8^-^, respectively) were analyzed for CD4 vs. CD47 expression. The red and turquoise circles mark the CD4^+^ subset and their unfractionated counterpart, respectively, that were used for the NGS experiment shown at the bottom of Fig. 2F.

We then administered Tamoxifen (TAM) via gastric gavage into adult MARC-11, and monitored Cre-mediated recombination in the peripheral blood and tail. The array was intact prior to TAM treatment (Fig. 2D, lane 1, 9). In contrast, by Day 2-8 post TAM (80 ug/g), the intact array had been recombined into shortened transgenes retaining predominantly zero or 1 gRNA gene (“zero-mer” or “monomer”), in addition to residual amounts of dimers (Fig. 2D, lane 2-8). At a higher dose of TAM (240 ug/g), zero-mer became more abundant and dimer fully vanished (lane 9-16). qPCR analysis confirmed that the proportions of the gRNA genes at P0 was reduced by TAM in a dose-dependent manner. For example, in the thymus, on Day 8 post TAM, the P0 gene abundance was reduced to 87% and 38% of the control level at 80 ug/g and 240 ug/g, respectively (Fig. 2E). Note that these experiments were done in F6 offspring (counting the founders as F1), indicating that the 11-gRNA repetitive sequence could be stably maintained for at least 6 generations without deletion.

We next used NGS to determine the representation of the various gRNA genes at P0 in the recombined array, finding it a function of their location on the array: the genes located toward the center were progressively underrepresented, resulting in a U-shaped curve (Fig. 2F, top). For example, for thymocytes, on Day 8 post TAM (80 ug/g), the gRNA gene at the beginning and the end of the 11-gRNA array (CD19 and CD4.5, respectively) constituted 56% and 8 % of the total P0 genes, respectively, whereas each of the three genes at the center (CD4.2, CD47.3 and CD4.3) ~ 2%. The 5 CD4 and 5 CD47 gRNA genes collectively constituted 20% and 14% of the total P0 genes, respectively, with the remaining 66 % of the reads represented by the single CD19 gRNA gene. The same trend was observed in other organs examined (Fig. 2F, top).

These data above indicate that on Day 8 post TAM, 87% of the thymocytes retained the P0 gRNA genes, of which 20% should express CD4 gRNAs and another 14% CD47 gRNAs. Thus, 17% and 12% of total thymocytes should express CD4 and CD47 gRNAs, respectively, with the two subsets mutually exclusive, and so should be the theoretical maximal proportions of the CD4^-^ or CD47^-^ subsets of the thymocytes. To test this idea, we used FACS to quantify CD4 and CD47 expression. In the thymus, CD4 is normally expressed in DP (TCR^-^D4^+^CD8^+^) and CD4 (TCR+CD4^+^CD8^-^) cells, while CD47 in both (Fig. 1G). To determine whether a fraction of these cells had lost CD4 or CD47 expression, we analyzed the two markers at the DP (TCR^-^D8^+^) and CD4 (TCR^+^CD8^-^) stages. Excitingly, as hoped for, cells lacking CD4 or CD47 marker were indeed detectable at both stages in a mutually exclusive manner, providing the first glimpse of the MARC technology at work. However, CD4 and CD47 expression was eliminated in only 5% of DP thymocytes (Fig. 2G, top middle), 3x lower than the theoretical maximum (17%), indicating that the efficiency of CD4 KO was quite moderate (~30%). The efficiency of CD47 KO was even less impressive, as only 1.4% of DP cells had lost CD47 expression, which was 9x lower than the theoretical maximum (12%; Fig. 2G, top middle). The KO efficiencies further decreases at the CD4 stage, where CD4^-^ or CD47^-^ cells became scarcer (Fig. 2G, bottom). Such low KO efficiencies were consistent with the poor U6 promoter activity in MARC-11 (Fig. 2C), but remarkably, did not seem to prevent *in vivo* screens, which provided the first piece of evidence for the inherent robustness of the MARC technology, as detailed below.

Genetic screens can be performed based on cellular abundance (“positive/negative selection”) or marker expression (“marker selection”)(Joung et al., 2017). We performed a marker selection screen using the CD4 molecule, by comparing P0 gRNA representations in the CD4^+^ thymocyte subset with that in the unfractionated counterpart comprising both CD4^+^ and CD4^-^ subsets (red and cyanine circles in Fig. 2G, respectively). Impressively, 4 out of 5 CD4 gRNA genes were apparently depleted in the CD4^+^ subset: CD4.1, CD4.2, CD4.3 and CD4.5, their abundance decreased by 23%, 13%, 50% and 13%, respectively (Fig. 2F, bottom), in broad agreement with the 30% overall KO efficiency mentioned above. These changes, albeit subtle, seemed specific, because the 5 CD47 gRNA genes were not depleted in the CD4^+^ subset; they were instead enriched, presumably as a consequence of the depletion of the 4 CD4 gRNA genes from the pool of the P0 gRNA genes. Of note, one of the 5 CD4 gRNA genes (CD4.4) was not depleted, perhaps because it was inactive; these gRNAs had been designed computationally and used without prior experimental validation.

In summary, the data above demonstrate that a transgene comprising 11 repeating units could be stably maintained for at least 6 generations *in vivo*; all the gRNAs on the array could be expressed following Cre-mediated recombination; and importantly, genetic screens seemed feasible despite the very poor gRNA expression. These data raised the exciting possibility that the basic elements of the MARC technology were all working. However, the gRNA expression apparently must be increased to make the tool practically useful.

### A 31-gRNA transgene inserted into the *H11* locus performed similarly to the 11-gRNA transgene

To corroborate and extend the conclusions based on MARC-11, we created a longer (31-gRNA) array, which was also inserted it into the *H11* locus but in a reverse orientation in the hope of increasing gRNA expression (Fig. 3A). The key component of the 31-gRNA array is a set of 8 gRNA genes, all designed to target the CD47 extracellular domain in order to facilitate their comparison (Fig. 3A, top). 4 of the 8 gRNAs (CD47.6-9) were newly designed and the remaining 4 (CD47.2-5) from the 11-array (CD47.1 was not selected because it targets a different CD47 domain). In addition to the 8 gRNA genes, the 31-gRNA array contains 22 gRNA genes (NC1-22) intended for use by dCas9-VP16, dCas9-KRAB or Cas13b, which are not expected to affect Cas9-mediated CD47 KO. These gRNA genes thus served as negative controls (NC) for the 8 CD47 gRNA genes.

**Fig. 3.**
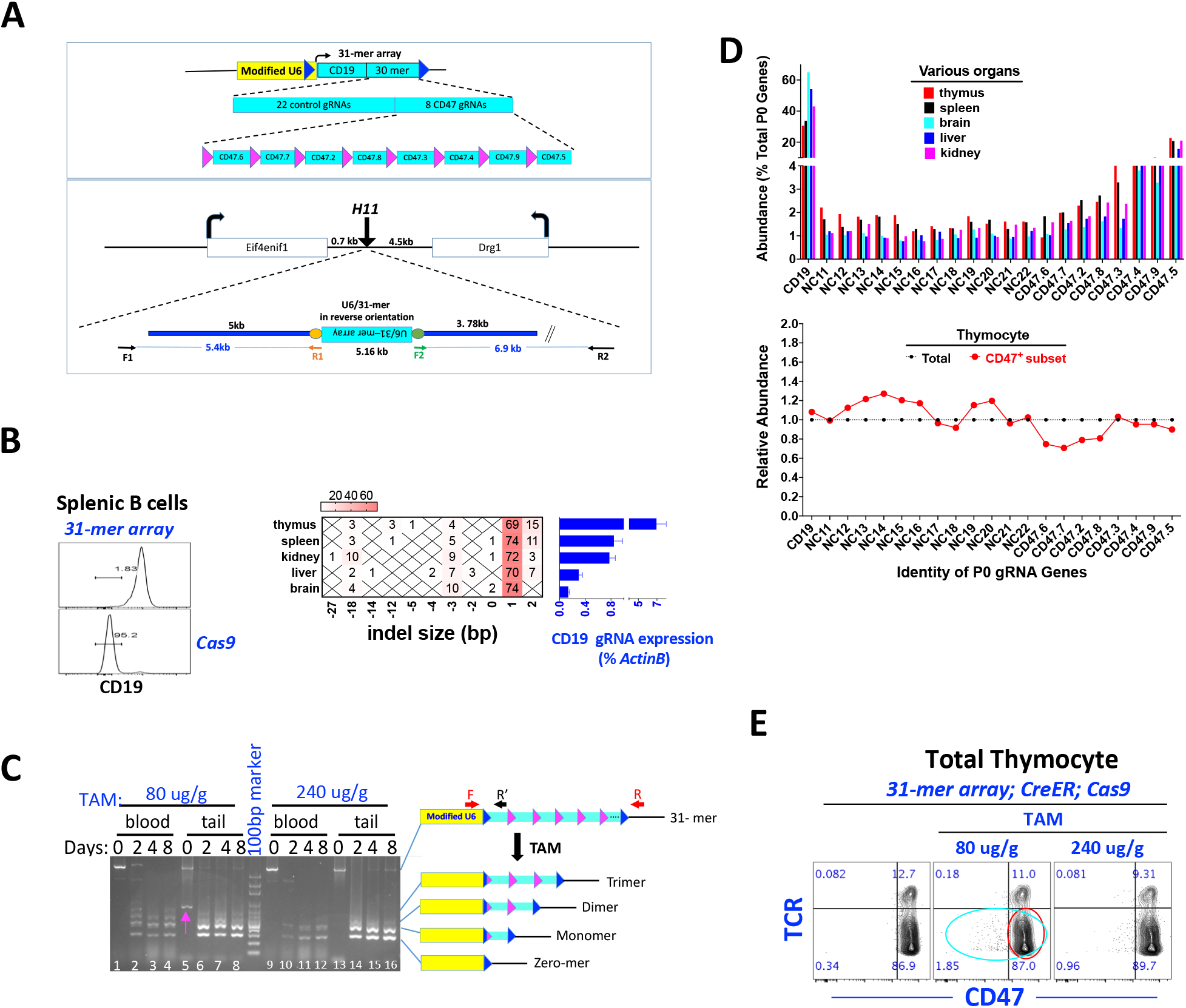
Creation and characterization of MARC-31. A) The 31-gRNA transgene, which is identical to the 11-gRNA transgene (Fig. 2A) except for the gRNA gene composition and transgene orientation relative to the *H11* locus. The founders were similarly screened and identified as in Fig. 2B (not shown). B-C) CD19 targeting (surface expression, indel formation and gRNA expression; Fig. 3B) and 31-gRNA transgene recombination (Fig. 3C), identical to Fig. 2 C-D except that MARC-31 instead of MARC-11 was analyzed. The pink arrow in C indicates a nonspecific amplicon. D) Representation of individual gRNA genes at P0 following recombination. Samples were harvested from F4 mice on Day 8 following TAM (80 ug/g) and analyzed by PCR as in Fig. 2F. Of note, the reverse primer R’ recognized the Cas9 scaffold but not the Cas13b scaffold, thus excluding the gRNA genes bearing the latter (NC1-10) from the analysis. The plot at the bottom compares P0 gRNA gene representation in the CD47+ subset of TCR^-^thymocytes relative to the unfractionated counterpart (see Fig. 3E for the populations). E) FACS analysis of CD47 vs. TCR expression in thymocytes. The red and turquoise circles mark the CD47+ subset and the unfractionated counterpart used for the NGS experiment shown at the bottom of Fig. 3D.

We found that in MARC-31, CD19 expression and the WT CD19 allele were nearly completely abolished, and the indels in various organs highly homogenous, suggesting very efficient KO in the early embryo (Fig. 3B, left and middle). However, in adult mice, CD19gRNA expression remained very low (<7% of *Actinb*; Fig. 3B, right). Importantly, as the 11-gRNA array, the 31-gRNA array was stably maintained (for at least 4 generations) and capable of dose-dependent recombination (Fig. 3C) to produce U-shaped representation of P0 gRNA genes (Fig. 3D, top). Consistent with the poor U6 promoter activity in adult MARC-31 mice, TAM could induce only minimal disruption of CD47 expression (Fig. 3E). Nevertheless, in the CD47+ subset of the thymocytes, the CD47 gRNA genes (except CD47.3) seemed depleted while the NC genes enriched (Fig. 3D, bottom), reminiscent of the reciprocal changes in the CD4 vs. CD47 gRNAs in CD4^+^ cells (Fig. 2F, bottom).

In sum, the 31-gRNA array was stably maintained and functional just as the 11-gRNA array, boosting our confidence in MARC, but the *H11* locus as a safe harbor was incompatible with high level gRNA expression, which was unexpected and disappointing.

### A 61-gRNA transgene randomly integrated using PiggyBac vector: stable maintenance with strong gRNA expression which enabled robust proof-of-concept *in vivo* screens

We next lengthened the 31-gRNA array to 61-gRNA by dimerizing the 30-gRNA array downstream of the CD19 gRNA gene, and inserted it into the genome randomly using a PiggyBac vector (Fig. 4A); we have decided against targeted insertion into a known safe harbor not only because the *H11* locus was incompatible with high-level gRNA expression, but the same seemed also true for the classic safe harbor *R26* locus (Fig. S1). Indeed, these “safe harbors” have been developed to support Pol II promoter function, whereas the U6 promoter, which is used by pol III, may need a divergent environment for optimal activity. To our knowledge, there is no well-defined safe harbor for the U6 promoter.

**Fig. 4.**
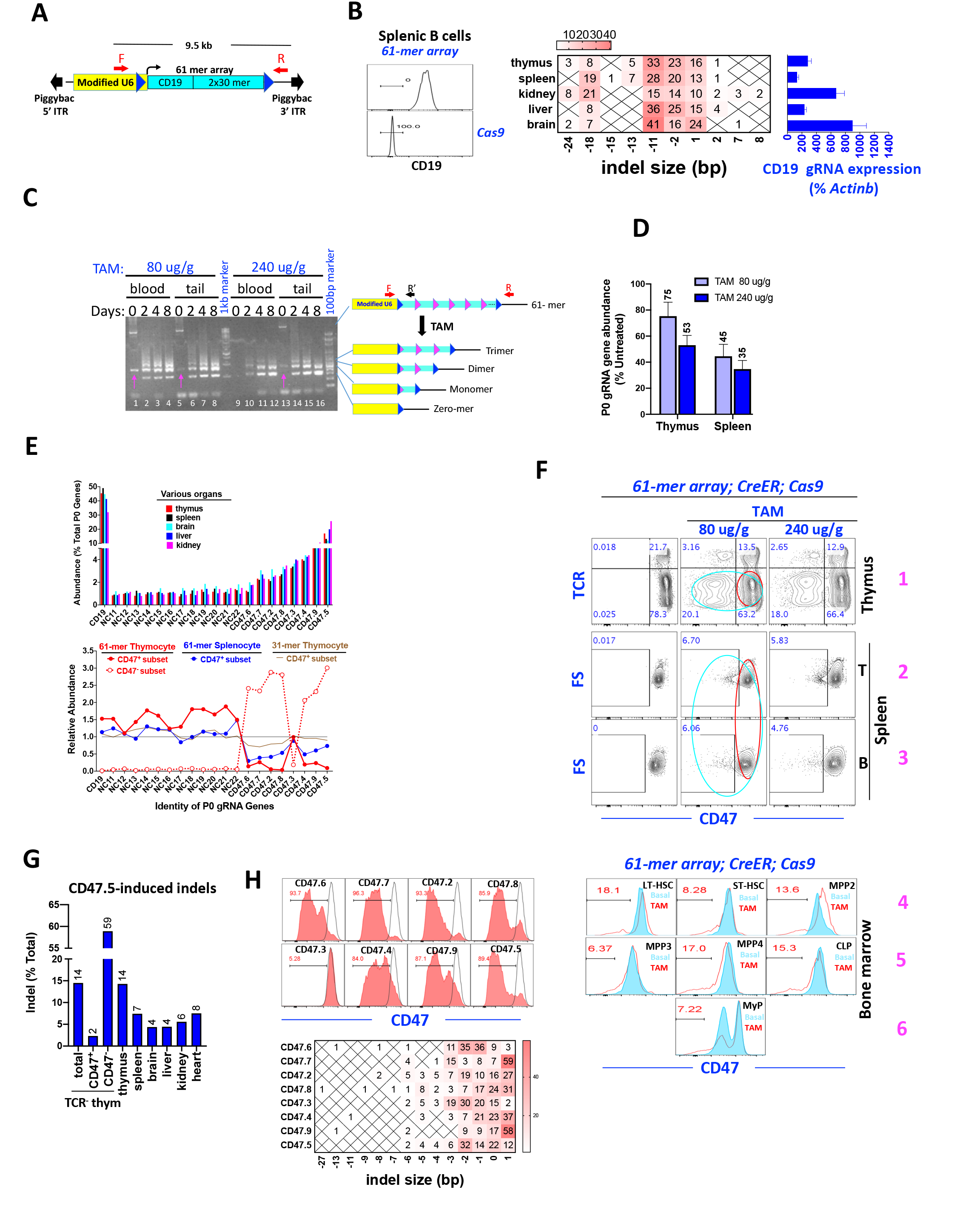
Creation and characterization of MARC-61. A) The 61-gRNA transgene. F/R, primer pair used for screening the founders, same as that used in Fig. 4C. B-D) CD19 targeting (surface expression, indel formation and gRNA expression; Fig. 4B), 61-gRNA transgene recombination (Fig. 4C) and the total abundance of various P0 gRNA genes as a whole on Day 8 post TAM (80 ug/g; Fig. 4D), identical to Fig. 2 C-F except MARC-61 instead of MARC-11 was analyzed. The pink arrow in C indicates nonspecific amplicons. The mice analyzed were of the F4 generation. E) Representation of individual gRNA genes at P0, analyzed as in Fig. 3D except that in the bottom plot, the CD47^-^ subset of TCR^-^ thymocytes (dotted red line) and CD47+ splenocytes (blue line) were included in addition to the CD47+ TCR^-^ thymocytes (solid red line; see Fig. 4F for the cell populations). The mice analyzed were of the F4 generation. For comparison, the bottom plot also displays the gRNA gene representation in the TCR^-^CD47+ thymocytes from MARC-31 (brown line, which is identical to the red line in Fig. 3D). F) FACS analysis of CD47 expression in the thymus (row 1), spleen (row 2-3) and bone marrow (row 4-6) from F4 mice. The red and turquoise circles mark the CD47+ subsets and their unfractionated counterparts used for the NGS experiment shown at the bottom of Fig. 4E. The gating strategy for various bone marrow cells is presented in Fig. S2. G) Proportions of the CD47 alleles bearing out-of-frame indels induced by the CD47.5 gRNA. The gRNA target site was amplified by PCR and analyzed by NGS. The mice analyzed were of the F4 generation. H) CD47 gRNA activity in a mouse cell line (N2a) expressing CD47. Plasmids expressing the gRNAs or Cas9 were co-transfected and the transfected cells drug selected before FACS analysis of CD47 expression on Day 8 post-transfection.

We obtained a total of 3 transgenic lines. Importantly, in the best line, the CD19 gRNA expression in adults was up to 9x higher than *Actinb* (150x of that in 11- or 31-gRNA mice), and this dramatically enhanced gRNA expression caused complete elimination of the CD19 protein and its WT allele (Fig. 4B).

The 61-gRNA transgene has so far been stably maintained for 4 generations. As expected, the array underwent dose-dependent recombination to give rise a U-shaped representation of the P0 gRNA genes as in MARC-31 (Fig. 4C-E), but importantly, the KO efficiency was much higher, consistent with the dramatic increase in the U6 promoter activity. Specifically, on Day 8 post TAM (80 ug/g), as much as 23% of thymocytes had lost CD47 expression (Fig. 4F, top). Since 75% of these thymocytes retained a P0 gRNA gene (Fig. 4D), of which 42% would express the CD47 gRNAs (Fig. 4E, top), the theoretical maximal proportion of the CD47^-^ thymocyte subset should be 31% of total thymocytes. This value slightly exceeded the observed value (23%), suggesting that CD47 was knocked out in 74% of thymocytes expressing CD47 gRNAs. Note that the efficiency of genome editing should be higher than 74%, as not all editing could produce null alleles. We conclude that the CD47 locus was efficiently (>74%) edited in the thymocytes in MARC-61.

In the spleen, the CD47^-^ subset constituted ~6 % of total splenocytes (Fig. 4F, middle and bottom rows), and 45% of the splenocytes retained a P0 gRNA gene (Fig. 4D), of which 37% would express CD47 gRNAs (Fig. 4E, top). The KO efficiency in the spleen was thus 35%, which was 2x lower than in the thymus, perhaps due to the relatively lower U6 promoter activity in the spleen (Fig. 4B, right).

As expected from the high KO efficiency in MARC-61 thymus, the CD47 gRNA genes (except CD47.3) were markedly depleted in the CD47+ subset of thymocytes while substantially enriched in the CD47^-^ subset (Fig. 4E, bottom, red lines). Note that these genes (except CD47.3) were also clearly depleted in the spleen, albeit to less extents than the thymus as predicted from the lower KO efficiency in the spleen (Fig. 4E, bottom, blue line). Thus, MARC-61 enabled robust genetic screens not only in the thymus with a high KO efficiency, and but in the spleen with a moderate KO efficiency, attesting to the robustness of the technology. Genetic screen should also be feasible in diverse cell types in the bone marrow, where the proportions of the CD47^-^ subsets (6%-18%) were comparable to the thymus and spleen (Fig. 4F, row 4-6).

We next examined the feasibility of genetic screens in solid organs (brain, liver, kidney and heart). As these organs are not readily amenable to FACS analysis, we quantified the indels at the CD47 locus as the surrogate for the loss of surface CD47 expression. We focused on the indels induced by last CD47 gRNA gene on the array (CD47.5), as this gRNA gene was the most represented among all the 8 CD47 gRNA genes (Fig. 4E, top). The CD47 genomic site targeted by CD47.5 gRNA was PCR-amplified and analyzed by NGS. As expected, the indels were substantially enriched and depleted in CD47^-^ and CD47^+^ subsets of thymocytes, respectively, and were more abundant in the thymus than the spleen (Fig. 4G). Importantly, the indels were readily detectable in all the solid organs, in agreement with their high U6 promoter activities, although the KO efficiencies did not strictly correlate with the promoter activities (Fig. 4B, right). Of note, the KO efficiencies in these organs were 2-3x lower than the thymus. As the mice were analyzed as early as Day 8 post TAM, prolonging the waiting period may increase the KO efficiencies. We conclude that genetic screens seem feasible in diverse organs in MARC-61.

As mentioned above, 7 out of 8 CD47 gRNA genes seemed to function well in mice. As expected, in a mouse cell line, in the presence of Cas9, these 7 gRNAs (but not the inactive CD47.3) eliminated CD47 expression (Fig. 4H top) and produced extensive indels at the target sites (Fig. 4H, bottom). Remarkably, CD47.3 was also able to efficiently edit its target site (Fig. 4H, bottom). Upon close examination, we discovered that the exon targeted by CD47.3 is annotated to be alternatively spliced and so its mutations may be inconsequential (not shown). These data, together with the fact that 4 out of 5 CD4 gRNAs seemed functional (Fig. 2F), suggest that 92% (12/13) of the computationally designed gRNAs proved effective at genome editing, which has implications for the design of the gRNA arrays (see further).

We conclude that the 61-gRNA array could be stably maintained for at least 4 generations, and in contrast to MARC-11/31, gRNA in MARC-61 was highly expressed, enabling efficient *in vivo* CD47 selection screens.

## Discussion

MARC rests on highly repetitive transgenes comprising tandemly linked gRNA genes. Repeat sequences are prone to silencing or deletion, which has been our major concern at MARC’s conception. It came as a pleasant surprise that the concern did not pan out. In particular, the 61- gRNA array has been maintained for 4 generations without any sign of deterioration. Furthermore, 2 different 100-gRNA arrays have also been created and have so far remained intact and functional for 2 generations (preliminary data). Thus, our gRNA arrays seem a fortunate exception to the general rule about the repeats. These lucky features, together with the ability of Cre to efficiently recombine the array, have materialized the MARC concept. While there is always room for optimization, MARC has emerged as a brand-new type of genetically engineered models with attractive potentials.

### MARC for *in vivo* genetic screens

MARC dramatically simplifies *in vivo* screening, thus promising to democratize it and unleash its full power: the PiggyBac constructs and transgenic mice can be made within weeks and months, respectively; the screening is done *in situ*, requiring no *ex vivo* manipulation but only routine procedures such as TAM administration and cell isolation; the key reagent (mice), once created, is self-perpetuating (via simple breeding); diverse cell types are targetable, including those inaccessible to viral libraries; the targeting can be global or spatiotemporal-specific depending on Cas9 expression patterns, offering flexibility in the experimental design; and finally, several distinct transgenes can be bred together and consolidated into a single mouse line, to reduce the screen scale (Peets et al., 2019) and more importantly, to enable combinatorial screens and hence the uncovering of genetic interactions (Sanson et al., 2019; Shen et al., 2017).

Compared with viral libraries, the capacity of MARC transgenes is low. This can be remedied using combinatorial screens: two and three distinct 100-gRNA transgenes used together are able to produce as many as 10^4^ and 10^6^ randomly multiplexed perturbations, respectively. More importantly, contrary to screening in cell lines, large-scale screening is often unfeasible in mice, either because the cells of interest are too scarce, or the “bandwidth” too narrow for the differentiating cells undergoing developmental transitions (e.g., only 500 gRNAs per mouse can be screened in the immune system reconstituted from gRNA-expressing bone marrow cells)(LaFleur et al., 2019). Under these conditions, *in vivo* screening can be scaled down using sub-libraries focused on special biological themes (such as kinases and epigenetic factors), as pointed out before (Doench, 2018). For such screens, MARC is readily applicable. For example, there are ~500 kinases in the mice and 167 epigenetic proteins frequently mutated in human cancers(Nanda et al., 2016), which could be covered by only 5 and 2 MARC-100, respectively. Indeed, it is our long-term plan to establish a comprehensive collection of MARC lines carrying various sub-libraries that together cover the entire genome, which should take only ~200 MARC-100 lines. This resource, complementary to the tens of thousands of single-gene KO lines produced by the International Mouse Phenotyping Consortium (Dickinson et al., 2016), may greatly facilitate mammalian functional genomics research.

### Other potential applications of MARC

First, MARC should enable cost-effective derivation of single-gene KO lines. The KO can be constitutive, using the null alleles from MARC. More importantly, the KO can be tissue-specific, using the recombined gRNA genes in addition to a transgene with tissue-specific Cas9 expression. Second, MARC can provide cells for single-cell RNA-Seq. This is analogous to “Perturb-Seq” where RNA-Seq is used to analyze the cultured cells transduced with a pooled lentiviral gRNA library (Dixit et al., 2016). However, MARC is culture/virus-free, and can offer diverse cell types for sequencing, including the cells hard to culture/transduce. Third, MARC can also be designed for other gRNA-based screens, such as gain-of-function screens using CRISPRa(Kampmann, 2018), or for RNA knockdown screens using Cas13 proteins(Abudayyeh et al., 2017; Cox et al., 2017; Konermann et al., 2018), the latter ideal for interrogating noncoding RNAs. Finally, MARC may be generally applicable to other multicellular organisms, thanks to the robustness and versatility of both the CRISPR-Cas and the Cre-Lox systems.

### Caveats

Given the low capacity of MARC, we recommend targeting each gene with a single gRNA rather than multiple ones as in conventional viral libraries. By necessity, this compact design will increase false negative rates in gene discovery, but perhaps only to a minor extent, as the majority of the gRNAs computationally designed are functional, as mentioned before. False positive rates should also rise due to off-target editing, but perhaps also only to a minor extent, as off-targets are largely avoidable using gRNA designing algorithms. Of note, if an gRNA indeed causes an interesting phenotype by fortuitously hitting an off-target, this could in fact lead to a lucky chance discovery. In any case, the error rates in the compact library can be further reduced using pre-validated gRNAs. In sum, although the “one gRNA per gene” design maximizes the library compactness only at some expense of robustness, the trade-off seems worthwhile.

Another shortcoming of MARC is the unequal representation of the P0 gRNA genes following recombination, with the gRNA genes at the ends of the array dominating the library whereas those in the middle underrepresented. This bias is tolerable for *in vivo* screens thanks to the power of targeted NGS, but will substantially increase the cost for single-cell RNA-Seq and the cost and labor for the derivation of single-gene KO lines. Preliminary data suggest replacing LoxKR3 with a novel LoxP variant might reduce the bias (not shown).

## Materials and Methods

### Plasmids and mice

Plasmid construction and sequences are detailed in Supplemental Information. To insert the 11- gRNA and 31-gRNA arrays into the mouse *H11* locus, the donor vectors carrying the arrays bearing homology arms were co-injected into the C57BL/6J zygotes with the Cas9 mRNA and the gRNA targeting the insertion site. The ensuing pups were screened with long-fragment PCR using the primer pairs F1/R1 and F2/R2 for the integration of the left and right homology arms, respectively (Figs. 2A-B, 3A). The PCR was done in PrimeStar GXL PCR reaction (TaKaRa, R050A). The primer sequences are:

F1: CTTGTGAGGGCCTACTGTGAC
R1: CTTTCCGGAGATAGGGTGTTA
F2: TTGCCCCTTTGTGTTCTCTTGTAG
R2: ATCGTGGGCATGTGACCTCTC

To insert the 61-gRNA array randomly into the mouse genome, the transposon carrying the array was co-injected into the zygotes with the mRNA encoding the PiggyBac transposase. The founders were identified by PCR and mated with WT mice to derive the lines bearing a singlecopy transgene. The CAG-Cas9 (JAX # 028555) and Ubc-CreER (JAX # #007179) transgenes were then introduced to generate the triple transgenic mice for subsequent analyses. The primers for genotyping are available upon request. Animal experiments were approved by the Animal Ethics Committee at ShanghaiTech, and performed in accordance with the institutional guidelines.

### TAM treatment

To prepare 50mg/ml stock solutions, 500mg TAM (Medchem Express, HY-13757A) was added to 9ml corn oil (Medchem express, HY-Y1888) before 1ml 100% EtOH was added. The solution was then incubated at 50°C for 30 min to dissolve TAM. TAM was delivered at ~5 ul (250 ug)/g or 1.7 ul (80 ug)/g via intragastric administration to adult mice.

### Transgene analysis following TAM treatment and *in vivo* screening

Cre-mediated recombination induced by TAM shortened the transgenes, and altered the abundance and identity of the P0 gRNA genes. To detect the shortening, we amplified both the intact and recombined products using Phanta Max (Vazyme P505) with primer pair F/R (Figs. 2D, 3C and 4C), which were then visualized by gel electrophoresis. ~50 ng genomic DNA, isolated from tail vein blood or tail tips, was used for PCR. The overall abundance of P0 gRNA genes was quantified with Luna^®^ Universal qPCR Master Mix (NEB, Cat: M3003S) using primer pair F/R’ (Fig. 2D) and 100 ng genomic DNA, with a fragment of *Brg* genomic DNA amplified as an internal control using the primer pair *Brg* F/R (Figs. 2E and 4D). To quantify the abundance of the P0 gRNA genes in various organs following TAM treatment (top panels in Figs. 2F, 3D and 4E), PCR was performed using AceTaq (Vazyme P412-03) on 200 ng genomic DNA with the primer pair F3/R3 (located at similar positions as F/R’) and the amplicons analyzed by NGS. To analyze the NGS data, we first used Cutadapt (2.4; https://cutadapt.readthedocs.io/en/stable/) to extract the spacer sequences from the reads. We then used Kallisto (v. 0.46.0; https://pachterlab.github.io/kallisto/manual) to map the extracted sequences to the MARC arrays. To do so, we first generated a kallisto index of gRNA arrays using the kallisto-index command with kmer set at 17. Using these kallisto indexes, we pseudoaligned the filtered fastq read files and quantitated the abundance of the aligned gRNAs using the following settings: kallisto quant-i MARC.index-t 5 fq. files.

For *in vivo* screening, the P0 gRNA gene representation in sorted cells were quantified by PCR using F3/R3 using 200-300 ng genomic DNA as described above. A critical parameter here is the amounts of the genomic DNA to be used in PCR, which dictates the fold of screen coverage. Following TAM (80 ug/g) treatment, the representations of the various gRNA genes at P0 were not even: those at the middle were the least abundant, constituting ~2% and ~1% of total P0 genes for MARC-11 and MARC-31/61, respectively (top panels in Figs. 2F, 3D and 4E). Besides, a fraction (20%-50%) of the cells lacked any gRNA genes at P0 due to over-recombination. Furthermore, for MARC-31/61, 1/3 of the gRNA genes, with the scaffold for Cas13b, could not be detected using F3/R3. Thus, in 200-300 ng genomic DNA, the least abundant P0 gRNA genes in MARC-11 and MARC-31/61 were represented by >300 and >100 cells, respectively. We found that under our PCR condition, a 60-fold screen coverage (using 40 ng MARC-11 genomic DNA) already sufficed for reproducible quantification of the P0 gRNA genes (Fig. S3). Indeed, in a previous RNAi-based *in vivo* screen, even 30-fold coverage of shRNA genes proved sufficient (Beronja et al., 2013). We have thus used 200-300 ng genomic DNA to ensure accurate quantification of gRNA representation in the *in vivo* screens.

The key primer sequences are:

F: TCCCCTGCCCCGGTTAATTTGCATA
R: CCGGCTCGTATGTTGTGTGGAATTGTGA
F: AGGCTTGGATTTCTATAAGAGA
R’: CCGACTCGGTGCCACT
F3: ACACTCTTTCCCTACACGACGCTCTTCCGATCTCACAAAAGGAAACTCACCCTAAC
R3: GTGACTGGAGTTCAGACGTGTGCTCTTCCGATCTGACTAGCCTTATTTAAACTTGCTAT
Brg F: CTGTATTTTGTCCGCATGATATACACT
Brg R: TGGCACTTCTTCAGGTTCTATGGG

### Analysis of indels at the CD19 and CD47 loci

We amplified ~900-bp and ~300-bp region encompassing the sites targeted by the CD19 and CD47.5 gRNAs, respectively, using 100-200 ng genomic DNA as templates, in Phanta Max (Vazyme P505). The primers used are listed below:

CD19 F: CTTTCCTCTATACGGGGACTGC
CD19 R: CAACTATGACTAACAGACAAGCGAGA
CD47 F: ACACTCTTTCCCTACACGACGCTCTTCCGATCTACCGAAGAAATGTTTGTGAAGT
CD47 R: GTGACTGGAGTTCAGACGTGTGCTCTTCCGATCTACAGTGGCAACCATATCTCAA

The amplicons were then subjected to Sanger sequencing (for CD19 indels; Figs 2C, 3B and 4C) or NGS (for CD47 indels; Fig. 4G). The Sanger sequencing data were analyzed by ICE (https://www.synthego.com/help/synthego-ice-analysis). The indels in the NGS data were detected essentially as described(Dow et al., 2015). Briefly, the sequence data were mapped to the mouse (MM10) genome using BWA version 0.7.10 (bwa mem-M). We then filtered the output SAM files for mapq ≥ 50 and retained only the reads containing a single indel.

### RNA extraction, qRT-PCR analysis of CD19 gRNA expression

Organs were harvested from adult triple transgenic mice prior to TAM treatment, snap-frozen in liquid nitrogen and homogenized in Trizol reagent. RNA was then extracted using Direct-Zol_RNA_Miniprep_Plus_Kit (Zymo Research, Cat: R2070) and reverse transcribed using HiScript^®^ II Q RT SuperMix (Vazyme, Cat: R223-01). CD19 gRNA expression in various organs was quantified using Luna^®^ Universal qPCR Master Mix (NEB, Cat: M3003S). The *Actinb* transcript served as an internal control. The primers were designed to span exons and are listed below.

gCD19 F: GAATGTCTCAGACCATATGGGGTT
gCD19 R: TGCCACTTTTTCAAGTTGATAACGG
Actin F: GGCTGTATTCCCCTCCATCG
Actin R: CCAGTTGGTAACAATGCCATGT

### Flow cytometry

Analytical FACS and electronic purification were performed using FACS Fortessa (BD Biosciences) and Aria III, respectively, and FACS data analyzed using Flowjo. Thymocytes and splenocytes from MARC-11 were stained with CD4, CD8, TCR, B220 and CD47 antibodies before the analysis of CD4 vs CD47 expression. For MARC-31/61, thymocytes and splenocytes were stained with TCR, B220 and CD47 antibodies before analyzing CD47 expression. Bone marrow cells from MARC-61 were stained with a cocktail comprising antibodies against lineage markers (Gr1, Ter119, CD3, B220, CD11b) and c-kit, Sca-1, IL-7Ra, Flt3, CD48, CD150 and CD47 antibodies before CD47 expression in various subsets of cells was analyzed. The details of the antibodies are provided in the Supplemental Information/

### Assessing CD47gRNA activities *in vitro*

The mouse neuroblastoma Neuro-2a (N2a; ATCC HTB-96) cells were cultured on 48-well plates at 37°C with 5% CO_2_ in DMEM (Life Technologies, 11965092) containing 10% FBS (HyClone, SH3008803), penicillin/streptomycin (Life, 15140122) and plasmocin prophylactic (Invivogene). Plasmid expressing Cas9 and blasticidin resistance gene (375ng) was co-transfected with a gRNA expressing vector (125ng) using Lipofectamine 3000 (Life Technologies, L3000008) following manufacturers’ instructions. 24h after transfection, blasticidin was added to 10μg/ml and cell culture continued for 7 more days before FACS analysis of CD47 expression.

### Contributions

TC designed the experiment and supervised the project with YC and JS. YC performed the experiments together with other co-authors.

## Acknowledgement

We thank Dr. Ruilin Sun at Shanghai Model Organisms Center, Inc for making the mouse lines and for providing the Rosa26-U6-gRNA mice, and Dr. Jun Lu for critical comments. The work is supported by the start-up funding from ShanghaiTech University to TC.

## Figure legends

**Fig. S1.**
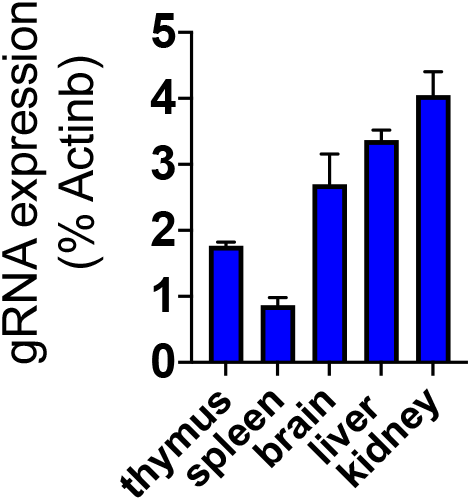
qRT-PCR analysis of gRNA expression from a mouse bearing U6-gRNA transgene inserted into the Rosa26 locus. Values are mean+/-SD from triplicate PCR.

**Fig. S2.**
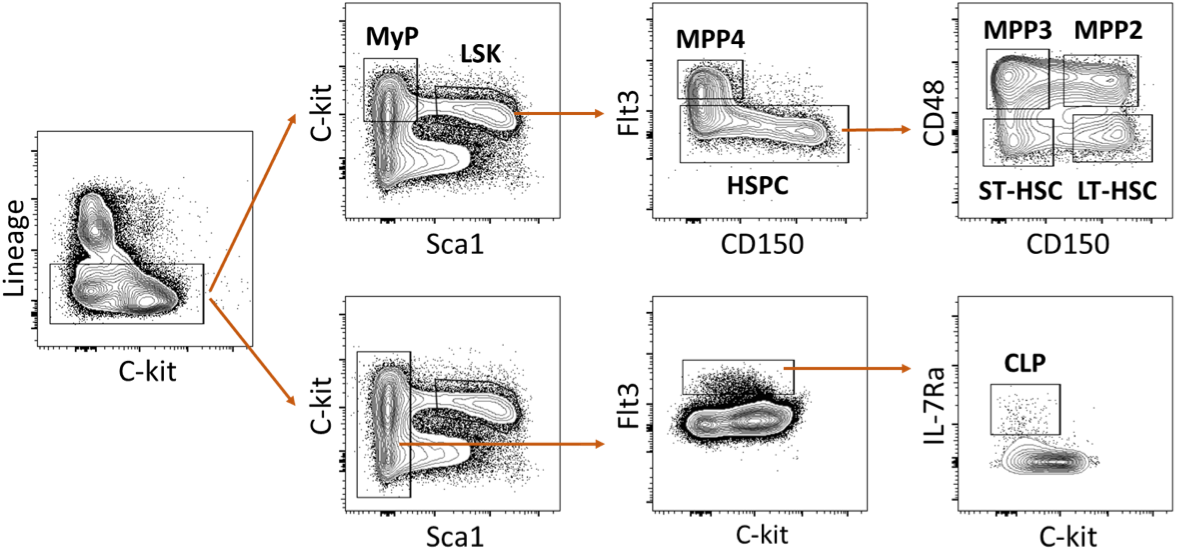
Gating strategy for the analysis of various subsets of bone marrow cells from MARC-61.

**Fig. S3.**
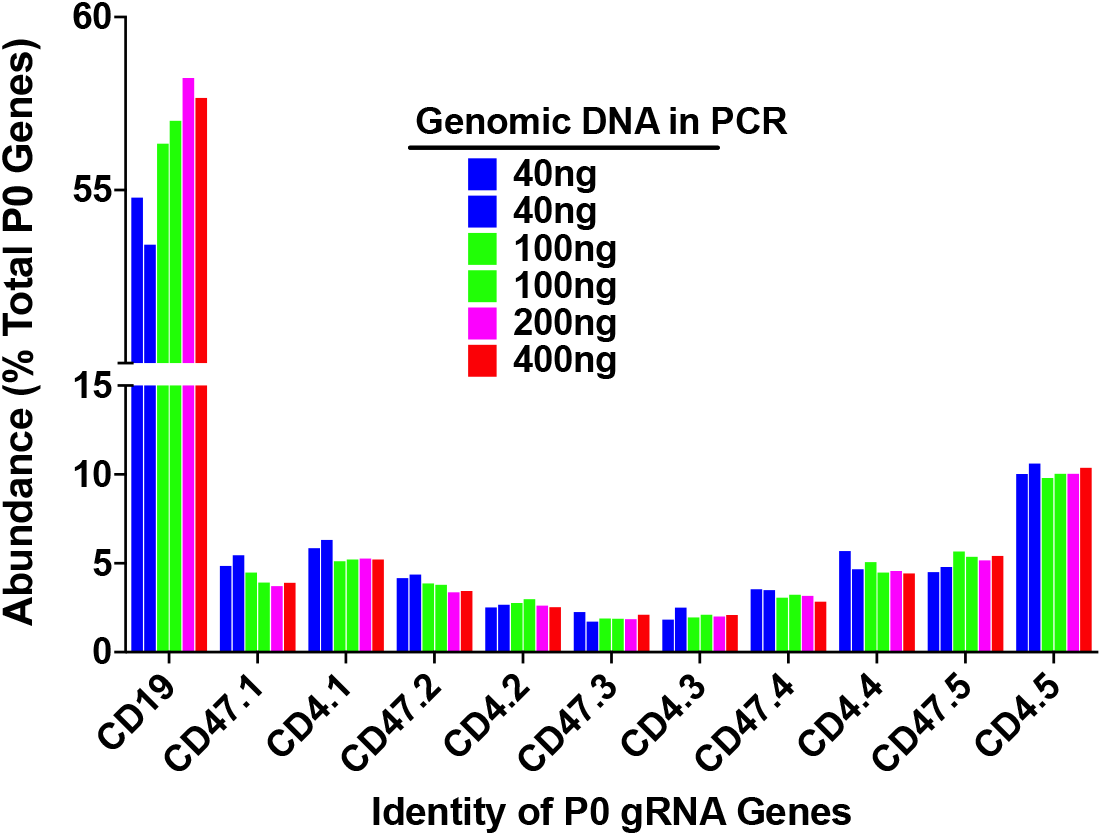
Determination of DNA doses required for accurate quantification of P0 gRNA gene representation. MARC-11 was treated with TAM (80 ug/g) and genomic DNA isolated from thymocytes.

## References

Abudayyeh, O.O., Gootenberg, J.S., Essletzbichler, P., Han, S., Joung, J., Belanto, J.J., Verdine, V., Cox, D.B.T., Kellner, M.J., Regev, A., et al. (2017). RNA targeting with CRISPR-Cas13. Nature 550, 280–284.

Araki, K., Okada, Y., Araki, M., and Yamamura, K. (2010). Comparative analysis of right element mutant lox sites on recombination efficiency in embryonic stem cells. BMC Biotechnol. 10, 29.

Beronja, S., Janki, P., Heller, E., Lien, W.-H., Keyes, B., Oshimori, N., and Fuchs, E. (2013). RNAi screens in mice identify physiological regulators of oncogenic growth. Nature 501, 185–190.

Bouabe, H., and Okkenhaug, K. (2013). Gene Targeting in Mice: a Review. Methods Mol. Biol. Clifton NJ 1064, 315–336.

Capecchi, M.R. (2005). Gene targeting in mice: functional analysis of the mammalian genome for the twenty-first century. Nat. Rev. Genet. 6, 507–512.

Chu, V.T., Graf, R., Wirtz, T., Weber, T., Favret, J., Li, X., Petsch, K., Tran, N.T., Sieweke, M.H., Berek, C., et al. (2016a). Efficient CRISPR-mediated mutagenesis in primary immune cells using CrispRGold and a C57BL/6 Cas9 transgenic mouse line. Proc. Natl. Acad. Sci. U. S. A. 113, 12514–12519.

Chu, V.T., Weber, T., Graf, R., Sommermann, T., Petsch, K., Sack, U., Volchkov, P., Rajewsky, K., and Kühn, R. (2016b). Efficient generation of Rosa26 knock-in mice using CRISPR/Cas9 in C57BL/6 zygotes. BMC Biotechnol. 16, 4.

Copeland, N.G., and Jenkins, N.A. (2010). Harnessing transposons for cancer gene discovery. Nat. Rev. Cancer 10, 696–706.

Cox, D.B.T., Gootenberg, J.S., Abudayyeh, O.O., Franklin, B., Kellner, M.J., Joung, J., and Zhang, F. (2017). RNA editing with CRISPR-Cas13. Science 358, 1019–1027.

Dickinson, M.E., Flenniken, A.M., Ji, X., Teboul, L., Wong, M.D., White, J.K., Meehan, T.F., Weninger, W.J., Westerberg, H., Adissu, H., et al. (2016). High-throughput discovery of novel developmental phenotypes. Nature 537, 508–514.

Dixit, A., Parnas, O., Li, B., Chen, J., Fulco, C.P., Jerby-Arnon, L., Marjanovic, N.D., Dionne, D., Burks, T., Raychowdhury, R., et al. (2016). Perturb-Seq: Dissecting Molecular Circuits with Scalable Single-Cell RNA Profiling of Pooled Genetic Screens. Cell 167, 1853–1866.e17.

Doench, J.G. (2018). Am I ready for CRISPR? A user’s guide to genetic screens. Nat. Rev. Genet. 19, 67–80.

Dong, M.B., Wang, G., Chow, R.D., Ye, L., Zhu, L., Dai, X., Park, J.J., Kim, H.R., Errami, Y., Guzman, C.D., et al. (2019). Systematic Immunotherapy Target Discovery Using Genome-Scale In Vivo CRISPR Screens in CD8 T Cells. Cell 178, 1189–1204.e23.

Dow, L.E., Fisher, J., O’Rourke, K.P., Muley, A., Kastenhuber, E.R., Livshits, G., Tschaharganeh, D.F., Socci, N.D., and Lowe, S.W. (2015). Inducible in vivo genome editing with CRISPR-Cas9. Nat. Biotechnol. 33, 390–394.

Friedrich, M.J., Bronner, I.F., Liu, P., Bradley, A., and Rad, R. (2019). PiggyBac Transposon-Based Insertional Mutagenesis in Mice. Methods Mol. Biol. Clifton NJ 1907, 171–183.

Guimont-Desrochers, F., Beauchamp, C., Chabot-Roy, G., Dugas, V., Hillhouse, E.E., Dusseault, J., Langlois, G., Gautier-Ethier, P., Darwiche, J., Sarfati, M., et al. (2009). Absence of CD47 in vivo influences thymic dendritic cell subset proportions but not negative selection of thymocytes. Int. Immunol. 21, 167–177.

Hippenmeyer, S., Youn, Y.H., Moon, H.M., Miyamichi, K., Zong, H., Wynshaw-Boris, A., and Luo, L. (2010). Genetic Mosaic Dissection of Lis1 and Ndel1 in Neuronal Migration. Neuron 68, 695–709.

Joung, J., Konermann, S., Gootenberg, J.S., Abudayyeh, O.O., Platt, R.J., Brigham, M.D., Sanjana, N.E., and Zhang, F. (2017). Genome-scale CRISPR–Cas9 knockout and transcriptional activation screening. Nat. Protoc. 12, 828–863.

Kampmann, M. (2018). CRISPRi and CRISPRa Screens in Mammalian Cells for Precision Biology and Medicine. ACS Chem. Biol. 13, 406–416.

Konermann, S., Lotfy, P., Brideau, N.J., Oki, J., Shokhirev, M.N., and Hsu, P.D. (2018). Transcriptome Engineering with RNA-Targeting Type VI-D CRISPR Effectors. Cell 173, 665–676.e14.

LaFleur, M.W., Nguyen, T.H., Coxe, M.A., Yates, K.B., Trombley, J.D., Weiss, S.A., Brown, F.D., Gillis, J.E., Coxe, D.J., Doench, J.G., et al. (2019). A CRISPR–Cas9 delivery system for in vivo screening of genes in the immune system. Nat. Commun. 10, 1–10.

Laurin, M., Gomez, N.C., Levorse, J., Sendoel, A., Sribour, M., and Fuchs, E. (2019). An RNAi screen unravels the complexities of Rho GTPase networks in skin morphogenesis. ELife 8.

Nanda, J.S., Kumar, R., and Raghava, G.P.S. (2016). dbEM: A database of epigenetic modifiers curated from cancerous and normal genomes. Sci. Rep. 6, 1–6.

Nguyen, D., and Xu, T. (2008). The expanding role of mouse genetics for understanding human biology and disease. Dis. Model. Mech. 1, 56–66.

Peets, E.M., Crepaldi, L., Zhou, Y., Allen, F., Elmentaite, R., Noell, G., Turner, G., Iyer, V., and Parts, L. (2019). Minimized double guide RNA libraries enable scale-limited CRISPR/Cas9 screens. BioRxiv 859652.

Rad, R., Rad, L., Wang, W., Cadinanos, J., Vassiliou, G., Rice, S., Campos, L.S., Yusa, K., Banerjee, R., Li, M.A., et al. (2010). PiggyBac Transposon Mutagenesis: A Tool for Cancer Gene Discovery in Mice. Science 330, 1104–1107.

Rahemtulla, A., Fung-Leung, W.P., Schilham, M.W., Kundig, T.M., Sambhara, S.R., Narendran, A., Arabian, A., Wakeham, A., Paige, C.J., Zinkernagel, R.M., et al. (1991). Normal development and function of CD8+ cells but markedly decreased helper cell activity in mice lacking CD4. Nature 353, 180–184.

Rickert, R.C., Rajewsky, K., and Roes, J. (1995). Impairment of T-cell-dependent B-cell responses and B-1 cell development in CD19-deficient mice. Nature 376, 352–355.

Rogers, Z.N., McFarland, C.D., Winters, I.P., Seoane, J.A., Brady, J.J., Yoon, S., Curtis, C., Petrov, D.A., and Winslow, M.M. (2018). Mapping the in vivo fitness landscape of lung adenocarcinoma tumor suppression in mice. Nat. Genet. 50, 483–486.

Rossant, J., and Spence, A. (1998). Chimeras and mosaics in mouse mutant analysis. Trends Genet. TIG 14, 358–363.

Ruzankina, Y., Pinzon-Guzman, C., Asare, A., Ong, T., Pontano, L., Cotsarelis, G., Zediak, V.P., Velez, M., Bhandoola, A., and Brown, E.J. (2007). Deletion of the developmentally essential gene ATR in adult mice leads to age-related phenotypes and stem cell loss. Cell Stem Cell 1, 113–126.

Sanson, K.R., DeWeirdt, P.C., Sangree, A.K., Hanna, R.E., Hegde, M., Teng, T., Borys, S.M., Strand, C., Joung, J.K., Kleinstiver, B.P., et al. (2019). Optimization of AsCas12a for combinatorial genetic screens in human cells. BioRxiv 747170.

Schramek, D., Sendoel, A., Segal, J.P., Beronja, S., Heller, E., Oristian, D., Reva, B., and Fuchs, E. (2014). Direct in Vivo RNAi Screen Unveils Myosin IIa as a Tumor Suppressor of Squamous Cell Carcinomas. Science 343, 309–313.

Shalem, O., Sanjana, N.E., and Zhang, F. (2015). High-throughput functional genomics using CRISPR-Cas9. Nat. Rev. Genet. 16, 299–311.

Shen, J.P., Zhao, D., Sasik, R., Luebeck, J., Birmingham, A., Bojorquez-Gomez, A., Licon, K., Klepper, K., Pekin, D., Beckett, A., et al. (2017). Combinatorial CRISPR-Cas9 screens for de novo mapping of genetic interactions. Nat. Methods 14, 573–576.

Ventura, A., Meissner, A., Dillon, C.P., McManus, M., Sharp, P.A., Van Parijs, L., Jaenisch, R., and Jacks, T. (2004). Cre-lox-regulated conditional RNA interference from transgenes. Proc. Natl. Acad. Sci. 101, 10380–10385.

Wang, G., Chow, R.D., Ye, L., Guzman, C.D., Dai, X., Dong, M.B., Zhang, F., Sharp, P.A., Platt, R.J., and Chen, S. (2018). Mapping a functional cancer genome atlas of tumor suppressors in mouse liver using AAV-CRISPR-mediated direct in vivo screening. Sci. Adv. 4, eaao5508.

Winters, I.P., Murray, C.W., and Winslow, M.M. (2018). Towards quantitative and multiplexed in vivo functional cancer genomics. Nat. Rev. Genet. 19, 741–755.

